# A high-resolution, country-wide scenario analysis of forest susceptibility to spruce bark beetle damage in Norway

**DOI:** 10.1101/2025.05.06.652523

**Authors:** Nicolas Cattaneo, Paal Krokene, Rasmus Astrup

## Abstract

The European spruce bark beetle (*Ips typographus* (L.)) poses a growing threat to Norway’s forests under climate change, particularly in extensive spruce-dominated forests. Because controlling active outbreaks is notoriously difficult, reducing forest susceptibility through proactive management is essential. In this study, we used high-resolution simulations covering all of Norway’s forested land (∼120,000 km^2^) to explore how different harvesting strategies affect long-term forest susceptibility to SBB damage. We compared a no-harvest baseline, a business-as-usual regime guided by economic criteria, and a risk-informed strategy that prioritizes harvesting of the most susceptible stands. Across two climate trajectories, we found that susceptibility-guided harvesting significantly reduced average forest susceptibility and fragmented high-risk areas, particularly in southern Norway. However, our results also reveal limitations to this approach: even with aggressive susceptibility-guided harvesting, long-term risk reductions plateau due to the homogenizing effects of even-aged forestry. Spatial connectivity analysis using electrical circuit theory further showed that susceptibility-guided harvesting disrupted landscape continuity among high-risk patches, suggesting a potential to constrain outbreak spread. These findings highlight the value of integrating susceptibility indicators and spatial connectivity metrics into forest planning to support adaptive, risk-informed management under climate uncertainty.

## Introduction

The European spruce bark beetle (*I. typographus* (L.)) is one of the most destructive pests in European forests (Anderegg et al., 2015). Under normal conditions, the spruce bark beetle (SBB) breeds in weakened or dying trees of its primary host Norway spruce (*Picea abies* (L.) H. Karst.). Such attacks introduce ecosystem heterogeneity by increasing light exposure and providing deadwood, thereby promoting biodiversity with minimal changes to the overall forest structure (Beudert et al., 2015). However, beetle populations can exceed a critical threshold and mass attack healthy trees following abiotic disturbances, such as prolonged drought, unusually warm temperatures or storms that reduce tree defenses and supply the beetles with suitable breeding material (Hlásny et al., 2019). The resulting outbreaks dramatically alter forest dynamics by triggering widespread tree mortality, depleting local carbon stocks, and reducing timber production. The altered habitat structure may, in turn, affect important ecosystem services and impact water quality, runoff, soil erosion, and nitrogen balance (Hlásny et al., 2019; Mikkelson et al., 2013).

In recent years, bark beetle outbreaks have intensified due to climate change and become more frequent in boreal and temperate conifer forests worldwide. In Europe, severe tree mortality events following SBB outbreaks have particularly affected Norway spruce monocultures in Central Europe, which are often planted outside the species’ natural distribution range (Hlásny et al., 2021). However, large volumes of spruce in northern Europe are also at risk, as climate change is expected to disproportionally affect northern areas (IPCC, 2021). Significant bark beetle damage is already observed in northern Europe, particularly in temperate spruce forests. For example, SBB killed approximately 34 million m^3^ of Norway spruce in southern Sweden following the exceptionally warm summer of 2018 (Wulff & Roberge, 2022). Further climate change will probably increase bark beetle damage also in the boreal spruce forests in Finland, Sweden, and Norway. Norway spruce is particularly abundant in Norway, accounting for approximately 44% of the country’s forest stock and around 70% of its annual timber harvests (SSB, 2025). As a result, large areas of Norway’s forests provide favourable habitat for the SBB, underlining the urgent need to develop effective strategies to mitigate bark beetle damage in the future.

Although abiotic factors play a significant role in predisposing spruce forests to bark beetle attacks, forest susceptibility is also strongly influenced by stand structure and forest management practices. Older stands with high volumes of Norway spruce are particularly vulnerable to infestation (Hlásny et al., 2019; Müller et al., 2022; Zimová et al., 2020). A strategy to mitigate this vulnerability is to limit the availability of suitable host trees (Seidl et al., 2016). The prevailing management approach in Norway’s production forestry relies on even-aged rotation systems, typically involving clear-cutting followed by replanting with the same species (Sevillano et al., 2025). Harvesting decisions are generally based on stand profitability and environmental regulations, while potential biotic or abiotic risk factors are not systematically considered. For example, harvesting tends to occur in stands classified as “ mature enough to be harvested,” and around 80% of harvesting takes place within 500 meters of a road, reflecting a strong emphasis on economic and logistical efficiency (Landbruksdirektoratet, 2023). However, increasing the resilience of future forests may require adjusting stand selection criteria to account for risk factors related to climate and pest outbreaks.

Implementing effective risk mitigation strategies typically requires a thorough quantitative understanding of how management actions impact not only the probability of a risk occurring but also the forest’s susceptibility to that risk (Hlásny et al., 2019). In this regard, new tools such as the SBB susceptibility index, developed by Nordkvist et al. (2023), have proven useful for analyzing and quantifying the impact of different management strategies on forest susceptibility to SBB damage (López-Andújar Fustel et al., 2024). Another element of risk mitigation is scenario simulation. This is a powerful approach for addressing the inherent uncertainty of the future, as it enables us to explore diverse possible outcomes. Scenario simulations envision alternative potential developments, allowing decision-makers to evaluate different strategies and prepare for a range of circumstances (Hoogstra-Klein et al., 2017). Using scenario simulations, we can test and compare how different forest management strategies influence the susceptibility of Norway spruce forests to SBB damage. Analyses of potential management impacts on forest structure at the landscape level can provide a comprehensive view of “ actionable space” and help identify opportunities and constraints for effectively addressing bark beetle risks. This could help forest managers anticipate risks, assess impacts, and develop adaptive management strategies in response to uncertain future conditions.

This study evaluates forest susceptibility to SBB damage under three contrasting management strategies: a conventional business-as-usual (BAU) approach, a risk-informed strategy that prioritizes the harvesting of highly susceptible stands, and a baseline no harvesting strategy. Using high-resolution simulations of forest susceptibility to SBB damage across all of Norway’s forested land, we examine long-term outcomes (2021–2100) under different climate trajectories. Our goal is to identify the actionable space that emerges when risk-related information is explicitly integrated into forest management, while still meeting future timber demands. By combining scenario analysis with decision-support tools, this study offers quantitative insights to support more resilient and adaptive forest planning under increasing disturbance pressure.

## Materials & Methods

### Study area and forest data

The study area included all forested regions of Norway, totaling ∼120 000 km^2^, or 37% of the country’s total land area. Climatic conditions vary much across this large area, with mean annual temperature ranging from −2 °C to around 6 °C and mean annual precipitation from 400 mm to 1300 mm. Norway’s forests are primarily composed of three dominant species: Norway spruce (*Picea abies* (L.) Karst.), Scots pine (*Pinus sylvestris* L.), and downy birch (*Betula pubescens*). Birch and other broadleaves constitute the largest share of the forest area (42%), while Norway spruce (42%) and Scots pine (30%) contribute most of the forest biomass (Breidenbach, Granhus, et al., 2020).

The forest information used in this study comes from the Norwegian Forest Resource Map SR16, which has a spatial resolution of 16 m × 16 m (Astrup et al., 2019). SR16 is based on empirical models fitted using the ca. 13 000 sample plots from the Norwegian National Forest Inventory (NFI) (Hauglin et al., 2021). The 16 m cell edge length was chosen as it is the closest integer that creates a square cell approximating the size of the circular NFI plots (250 m^2^). (Astrup et al., 2019). SR16 consists of a series of raster maps describing various forest state variables that can be used to initialize forest growth simulations. The main SR16 variables used in this study include the site index (SI = the average height of the 100 thickest trees per hectare at a reference age of 40 years, as defined by Tveite & Braastad, (1981)), mean stand age (in years), basal area (BA, in m^2^/ha), number of trees (N, stems per hectare), dominant stand height (H, in meters), and standing volume with bark (V, in m^3^/ha). Additionally, we used a raster map indicating the dominant tree species or group for each cell (Norway spruce, Scots pine, or broadleaves) (Breidenbach, Waser, et al., 2020). SR16 also provides stand-like aggregations of pixels (Astrup et al., 2019), which were used when stand-level calculations were required.

Additional in-house spatial datasets required to simulate forest management decisions were also used, including information on harvesting costs (e.g., digital terrain models, proximity to roads, and environmentally protected areas). Timber demands at the country level under the representative concentration pathway RCP4.5 climate trajectory was derived from the GLOBIOM-forest model (Lauri et al., 2021). This model is based on the Nationally Determined Contribution scenario and further details are provided in Vergarechea et al. (2023) and Blattert et al. (2023). For the near present-day climate (NPC) trajectory (see below), future timber demand was derived by keeping the RCP4.5 timber demand for the years 2021–2025 constant throughout the simulation.

### Forest projections and climate scenarios

Forest projections were generated using the PixSim forest growth simulator (Cattaneo et al., 2024), which has been previously used for countrywide and regional analyses of the impact of climate and forest management on surface albedo (Bright et al., 2024) and streamflow (Huang et al., 2025). The simulator is designed to operate at the pixel level of high-resolution, wall-to-wall forest resource maps like SR16. It efficiently implements stand-level forest growth models for each pixel, enabling rapid large-scale growth simulations. The simulator includes a module that incorporates climate effects on forest growth and a module that implements forest management decisions. In this study, all simulations were initialized with near-present-day forest characteristics using 2021 as the reference year. Forest growth over time was simulated in 5-year time steps using the species-specific empirical stand-level growth models developed by Maleki et al. (2022).

To simulate forest projections, we built six scenarios by combining two climate trajectories with three forest management regimes (described below). The climate trajectories included one where the climate changes under a moderate mitigation scenario, as given by RCP4.5, and another where the climate is kept constant near current conditions (NPC). The NPC climate trajectory provides a baseline to evaluate effects of climate change on forest susceptibility to bark beetle damage. Key climate variable inputs covering the period 1971–2100 were obtained from the Norwegian Meteorological Institute (including daily mean temperatures and precipitation). The future climate trajectory following RCP4.5 originated from a combination of three regional climate model simulations from the EURO-CORDEX archive (Wong et al., 2016). Data was first downscaled to a 1 × 1 km grid and bias corrected and then further downscaled to the 16 m grid used in SR16. Methodological details on the downscaling to the 16 m grid can be found in Bright et al. (2024). Climate change feedbacks on forest growth over time were incorporated through changes to the site index, using the climate-sensitive site index models of Antón-Fernández et al. (2016).

### Forest management regimes

We simulated three contrasting forest management regimes to evaluate how susceptibility to spruce bark beetle (SBB) damage developed under different harvesting strategies, with the idea of estimating the maximum susceptibility reduction that can be achieved when harvest allocation is guided exclusively by stand-level susceptibility values. The first regime, no harvesting (NoH), excludes any harvesting activity, allowing forests to grow undisturbed throughout the simulation period. NoH serves as a baseline to assess how harvesting interventions affect long-term susceptibility. The second regime, business-as-usual (BAU), reflects the current dominant management approach in Norway, involving even-aged rotation forestry based on national guidelines, where harvesting is guided by stand profitability and environmental constraints. BAU practices include clear-cutting, replanting, and maintaining existing species composition. On average, about 0.45% of the forest area in Norway is harvested each year under this regime (Norwegian Ministry of Climate and Environment, 2020), indicating a relatively low level of management intensity. The third regime, susceptibility-guided harvesting (BBprior), takes a radically different approach: stands are prioritized for harvesting solely based on their mean susceptibility to SBB. All stands in Norway are ranked in descending order of susceptibility and harvesting proceeds from the most susceptible downward, until national timber demand is met for each timestep. Under BBprior all harvested stands are replanted with the same species.

### Spruce bark beetle susceptibility

We quantified forest susceptibility to SBB attack using the SBB susceptibility index developed by Nordkvist et al. (2023). The index provides a relative measure of forest predisposition to SBB attack, rather than a direct probability of damage, and has been used in previous studies to evaluate the impact of different management strategies on forest susceptibility to SBB damage (López-Andújar Fustel et al., 2024). The index incorporates several key parameters that influence forest susceptibility to SBB damage, such as stand characteristics (e.g., Norway spruce volume, mean stem diameter, stem density, age structure, and birch volume), soil moisture, and temperature sum for the growing season. The SSB susceptibility index ranges from 0 to 3.66, with higher values indicating greater susceptibility. Three initial conditions must be met for the index to exceed zero: a temperature sum > 745 degree days, a Norway spruce volume > 0, and a stand mean stem diameter of Norway spruce ≥ 20 cm. The contribution of each parameter to the total index value is weighted according to its relative importance in increasing susceptibility, as derived from scientific evidence (Nordkvist et al., 2023). For instance, Norway spruce volume and temperature sum are considered highly influential and are each assigned a weight of 1. Age structure has a moderate weight of 0.5 due to ambiguous evidence. Birch volume is assigned a weight of 0.2, reflecting inconsistent and sometimes contradictory empirical evidence. Norway spruce-dominated stands have a susceptibility value of zero until the average stem diameter reaches 20 cm (because the beetles usually do not colonize trees that are smaller than this). Thus, during this phase—between harvesting and mean stand diameter reaching the 20 cm threshold—stands are considered to be non-susceptible. We computed the SBB susceptibility index at the beginning of each 5-year simulation step. The index was calculated only for pixels located in stands with at least one spruce-dominated pixel and a spruce volume > 0. Daily mean temperatures for the years 2016–2100 were used to compute the temperature sum for the growing season (further details on parameter calculations are in the Appendix).

To compare levels of SBB susceptibility across the six scenarios, we analyzed the temporal development (2021-2100) of mean susceptibility at the whole-country level for each scenario. The following procedure was used to calculate the country-level mean susceptibility for each 5-year simulation time-step. First, we identified the most common tree species in each stand (by counting the number of pixels dominated by each tree species) and focused only on stands where spruce was the dominant species —specifically, those in which more than 50% of the pixels were classified as spruce-dominated. Second, using all the pixels within these stands, we calculated a country-level mean susceptibility index. Pixels with zero susceptibility were excluded from the calculations and instead used to determine the area of non-susceptible spruce-dominated forest. To further investigate the mean susceptibility values, we plotted the distribution of spruce volume across the observed values of the SBB susceptibility index. Total standing volume for each scenario and simulation time step was broken down into 0.1-class intervals based on the observed SBB susceptibility index, and the temporal development of the distributions was analyzed.

### Landscape connectivity

Finally, to address spatial patterns of SBB susceptibility across scenarios, we performed a large-scale connectivity analysis using a model based on electrical circuit theory (McRae et al. 2008), implemented in the Circuitscape software (Anantharaman et al. 2020). The main input for modeling connectivity using Circuitscape is a resistance surface, which represents landscape features based on the degree to which they impede movement. To create the resistance surface, we generated a 500×500 m resolution map of maximum SBB susceptibility, where each 500×500 m cell was assigned the index value of the 16×16 m pixel inside it that had the highest value. The resistance map covered a 79,294 km^2^ area in southern Norway that included most of Norway’s spruce-dominated forests (Breidenbach, Granhus, et al., 2020; Breidenbach, Waser, et al., 2020). The 500 m resolution was selected to address computational limitations and because bark beetle infestation spread is strongly distance-dependent, with 95% of new SBB infestations typically occurring within 500 m of the previous year’s infestations (Kautz et al., 2011). The resistance surface was created by taking the maximum SBB susceptibility index value found within each 500×500 m tile and subtracting this value from 3.66 (the maximum possible index value). This transformation reversed the scale of the SBB susceptibility index, ensuring that areas with higher susceptibility corresponded to lower resistance to SBB movement. To ensure greater contrast in areas where the SBB susceptibility index was zero, these pixels were assigned a resistance value of 100. The connectivity metric we used was the mean pairwise effective resistance (MPER), which quantifies overall network connectivity. MPER is a measure of the cost of traveling across the landscape, where higher MPER values indicate poorer connectivity between patches with high susceptibility, while lower MPER values reflect more potential pathways for movement and higher overall connectivity (O’Brien et al., 2023). MPER was calculated as the average of all pairwise effective resistances between 75 focal nodes that were evenly distributed along the perimeter of the 79,294 km^2^ area of interest. We used the same number and fixed locations of focal nodes for all scenarios to facilitate comparisons. MPER was computed for every simulation time step of each scenario, and we analyzed its temporal development to examine how different harvesting strategies affected the connectivity between areas with high susceptibility to SBB damage.

## Results

The temporal development of forest susceptibility to SBB at the country level revealed distinct differences across the three management regimes (NoH, BAU, BBprior) and two climate trajectories (RCP4.5, NPC; Figure 1). BBprior consistently yielded the lowest SBB susceptibility values, demonstrating the effectiveness of this risk-informed management regime in reducing forest susceptibility to SBB damage (Figure 2). This reduction was more pronounced under the NPC climate trajectory compared to RCP4.5, particularly during the first 40 years of the simulations (Figure 1). In contrast, the NoH regime resulted in the highest SBB susceptibility values over time, as no harvesting led to an accumulation of older stands, which are typically associated with higher susceptibility. BAU showed similar, but slightly lower, SBB susceptibility values compared to NoH. For both these management regimes there was a general trend of increasing susceptibility over time under both climate trajectories (Figure 1). Under BBprior, susceptibility decreased sharply in the first 20 years of the simulations, before leveling off or increasing towards 2090. Warmer conditions under RCP4.5 amplified risks, leading to higher SBB susceptibility values, while leaving the temporal trends associated with the different management regimes largely unchanged. For BBprior, the RCP4.5 climate trajectory increased susceptibility by up to 2.34% compared to NPC; however, these differences were reversed by the end of the planning horizon (2100).

**Figure 1.**
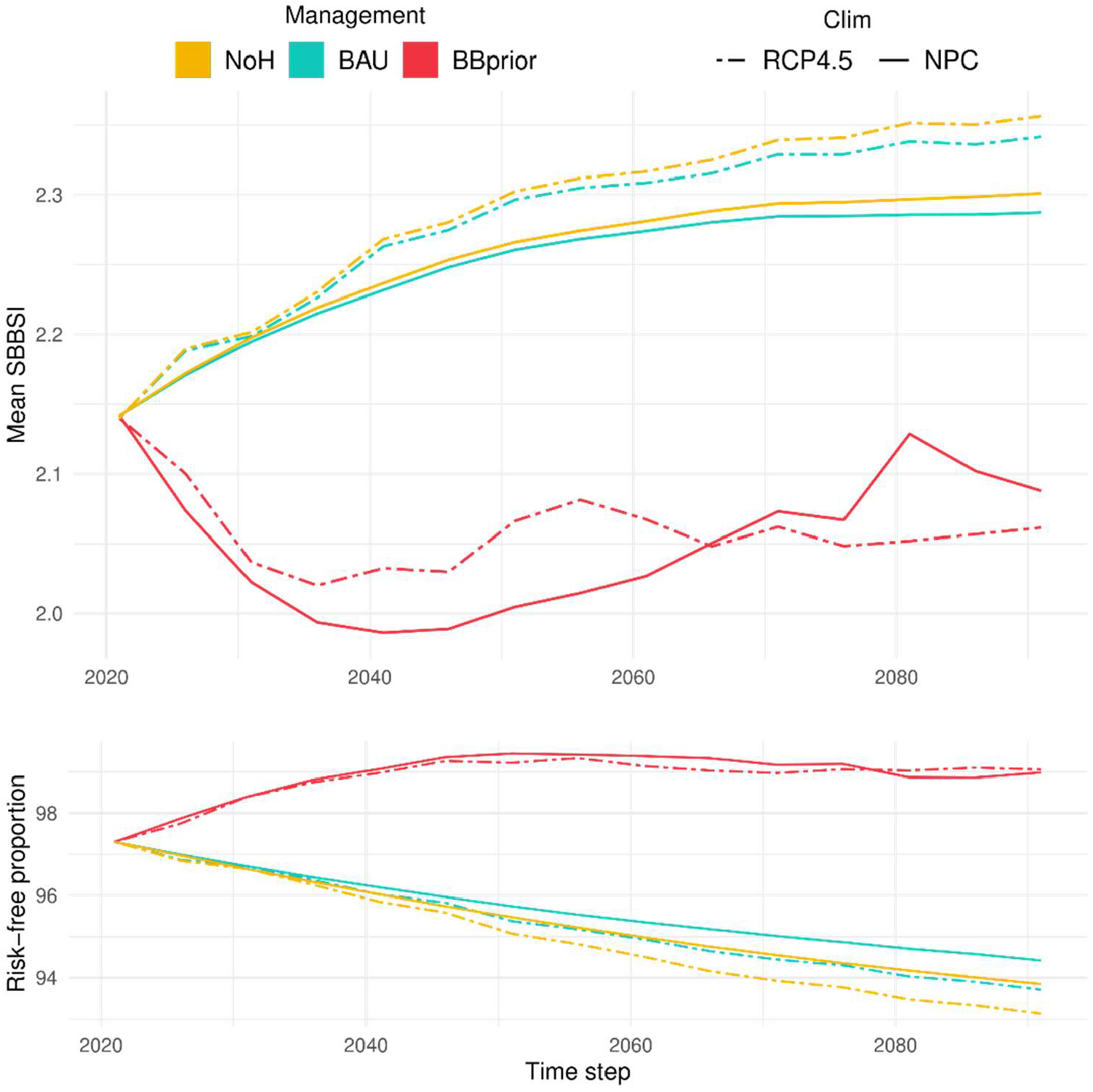
Temporal development of mean spruce bark beetle susceptibility index (SBBSI) (upper panel) and non-susceptible proportion of spruce-dominated forests (lower panel) at the whole-country level under different management regimes and climatic trajectories. Simulated management regimes were: no harvesting (NoH), business as usual (BAU), and prioritized harvesting in stands with high SBBSI (BBprior). Two different climate trajectories were considered: Representative Concentration Pathway 4.5 (RCP4.5) (dashed lines) and near present-day climate (NPC) (solid lines).

**Figure 2.**
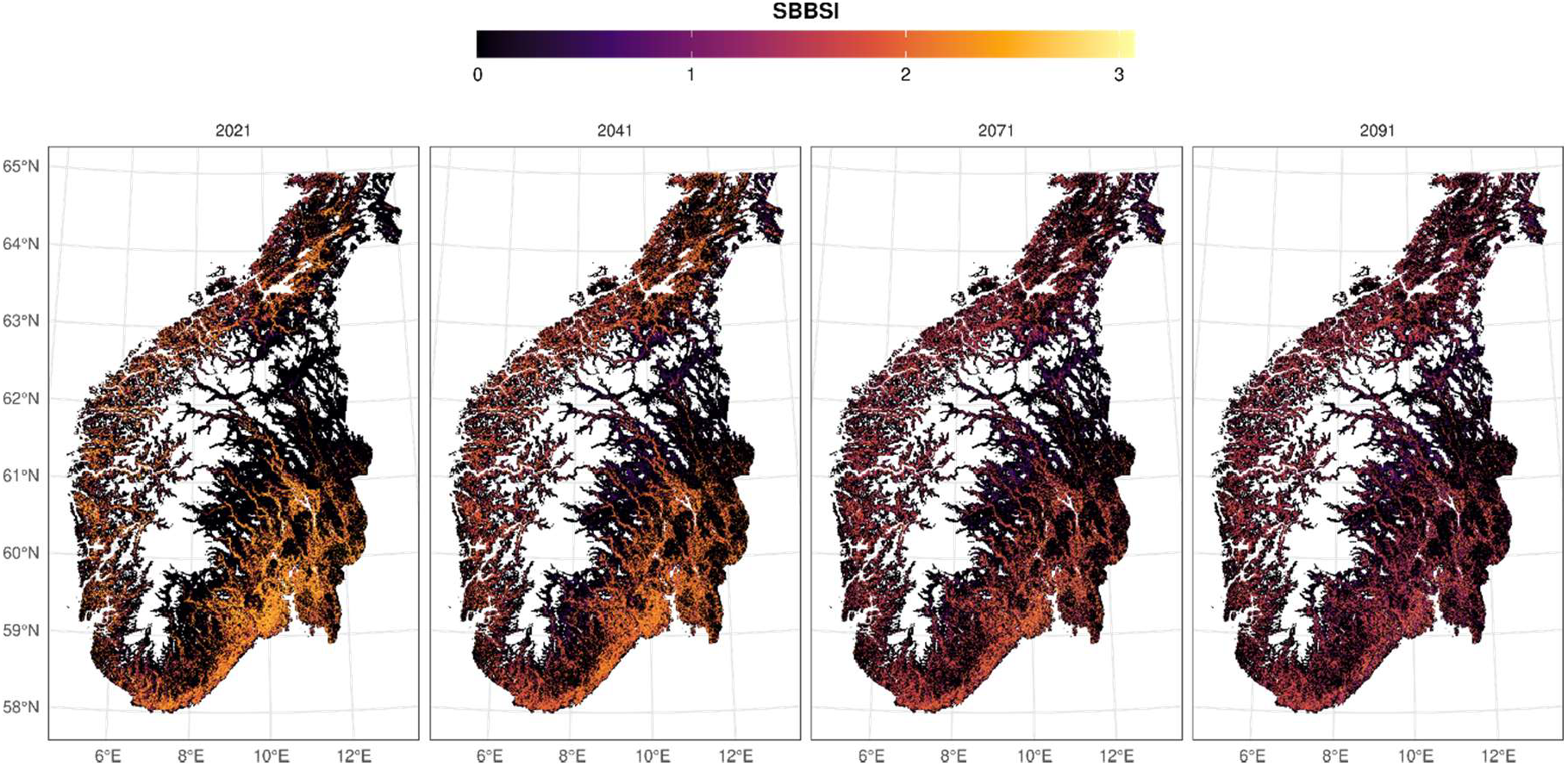
Temporal development of forest susceptibility to spruce bark beetle (SBB) damage across southern Norway. The maps correspond to four time steps from the simulation scenario combining Representative Concentration Pathway 4.5 (RCP4.5) and prioritizing the harvest of stands with the highest susceptibility (BBprior). For better visualization, we plotted maximum SBB susceptibility index (SBBSI) values at a resolution of 500×500 m. Country-level mean SBBSI values are 2.14, 2.03, 2.06, and 2.06 for the time steps 2021, 2041, 2071, and 2091, respectively.

The temporal development of non-susceptible spruce-dominated forest area at the country level reflected the temporal changes in mean SBB susceptibility index (Figure 1). Management regimes that led to increased susceptibility over time (NoH, BAU) showed a corresponding decline in non-susceptible area, while the BBprior regime which decreased susceptibility led to an increase in non-susceptible forest area for the first 20 years of the simulations. Differences in non-susceptible area between climate trajectories were less pronounced compared to those observed for mean SBB susceptibility (Figure 1).

The distribution of standing spruce volume across SBB susceptibility classes highlighted the effects of management and climate trajectories on susceptibility (Figure 3). Under BBprior, total spruce volume with high SBB susceptibility decreased substantially over time and shifted towards lower susceptibility classes. This indicated the success of this management regime in reducing both mean SBB susceptibility at the national level and the total amount of timber at risk. In contrast, NoH maintained higher volumes of spruce with high SBB susceptibility due to the absence of intervention. This trend intensified over time as the spruce volume in older, high-risk stands accumulate. Similarly, the BAU regime lead to a concentration of volume in highly susceptible stands. However, harvesting activities distinguished BAU from NoH, particularly in the highest susceptibility classes. As seen in Figure 3, warmer temperatures in RCP4.5 accelerated shifts in susceptibility, exposing spruce volume to risk much faster than under NPC. Towards the end of the simulation (2091), most spruce volume was concentrated in the two highest risk classes under RCP4.5, whereas NPC retained a substantial volume in intermediate-risk classes.

**Figure 3.**
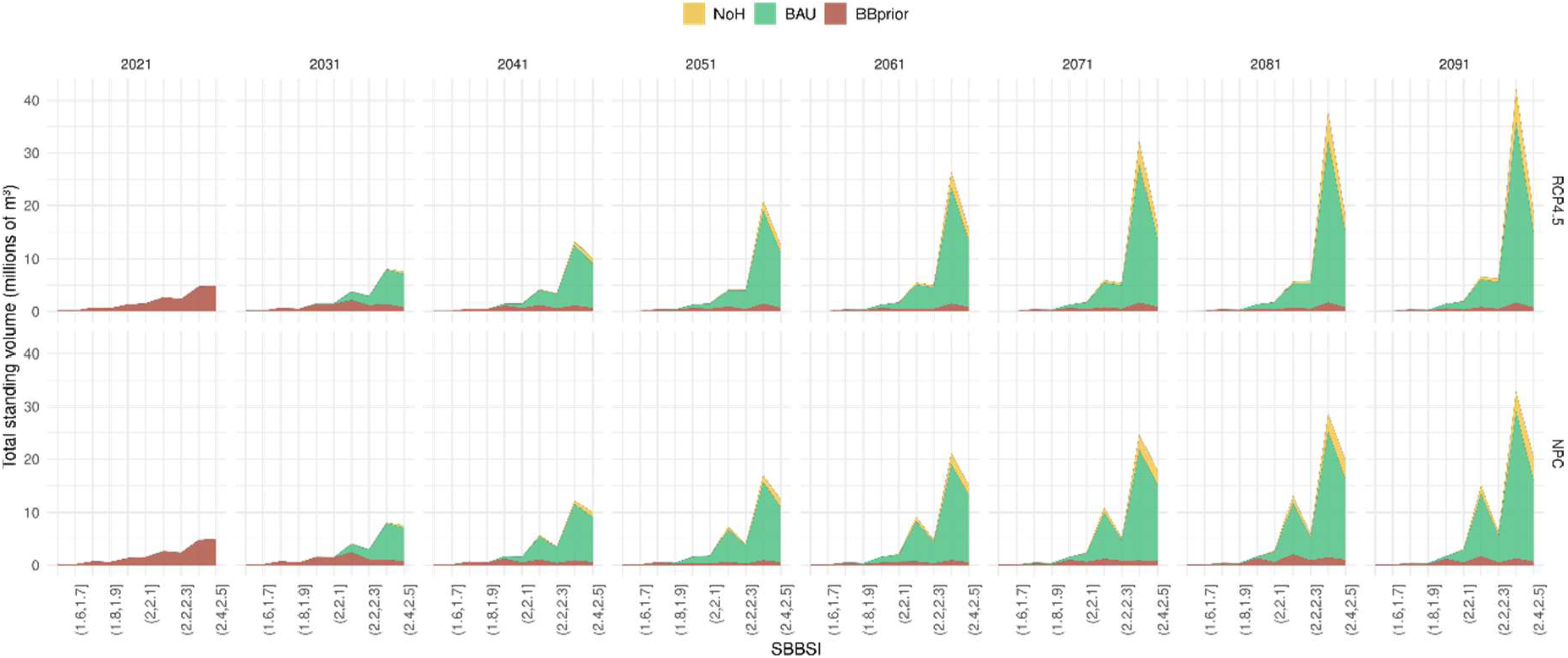
Temporal development of total standing volume of Norway spruce with different susceptibility to spruce bark beetle (SBB) damage under two climate trajectories: Representative Concentration Pathway 4.5 (RCP4.5) and near present-day climate (NPC). Development is shown for three forest management regimes: no harvesting (NoH), business as usual (BaU), and prioritize harvesting in stands with high susceptibility (BBprior). SBBSI: SBB susceptibility index.

Since climate change only altered the magnitude of the SBB susceptibility index without substantially changing its temporal dynamics, we present the results of the connectivity analysis only for the RCP4.5 scenario. While the overall connectivity between high-susceptibility areas, as expressed by MPER, varied between management regimes throughout the simulation period (Figure 4), differences between NoH and BAU remained minimal. In general, harvesting, regardless of the management regime, consistently reduced MPER over time, increasing connectivity between high-susceptibility areas toward the end of the simulations. However, the BBprior management regime showed consistently higher MPER values compared to BAU throughout almost the entire simulation, indicating lower connectivity between susceptible areas. The differences in MPER were evident from the second simulation step, as soon as harvesting began, and persisted throughout the study period. The temporal development of MPER suggests that the spatial distribution of susceptible areas changed over time, with notable differences between management regimes in the connectivity of high-risk forest patches. Figure 5 presents a cumulative current density (CCD) map for the BBprior management regime, visualizing current flows across the landscape, highlighting connectivity patterns.

**Figure 4.**
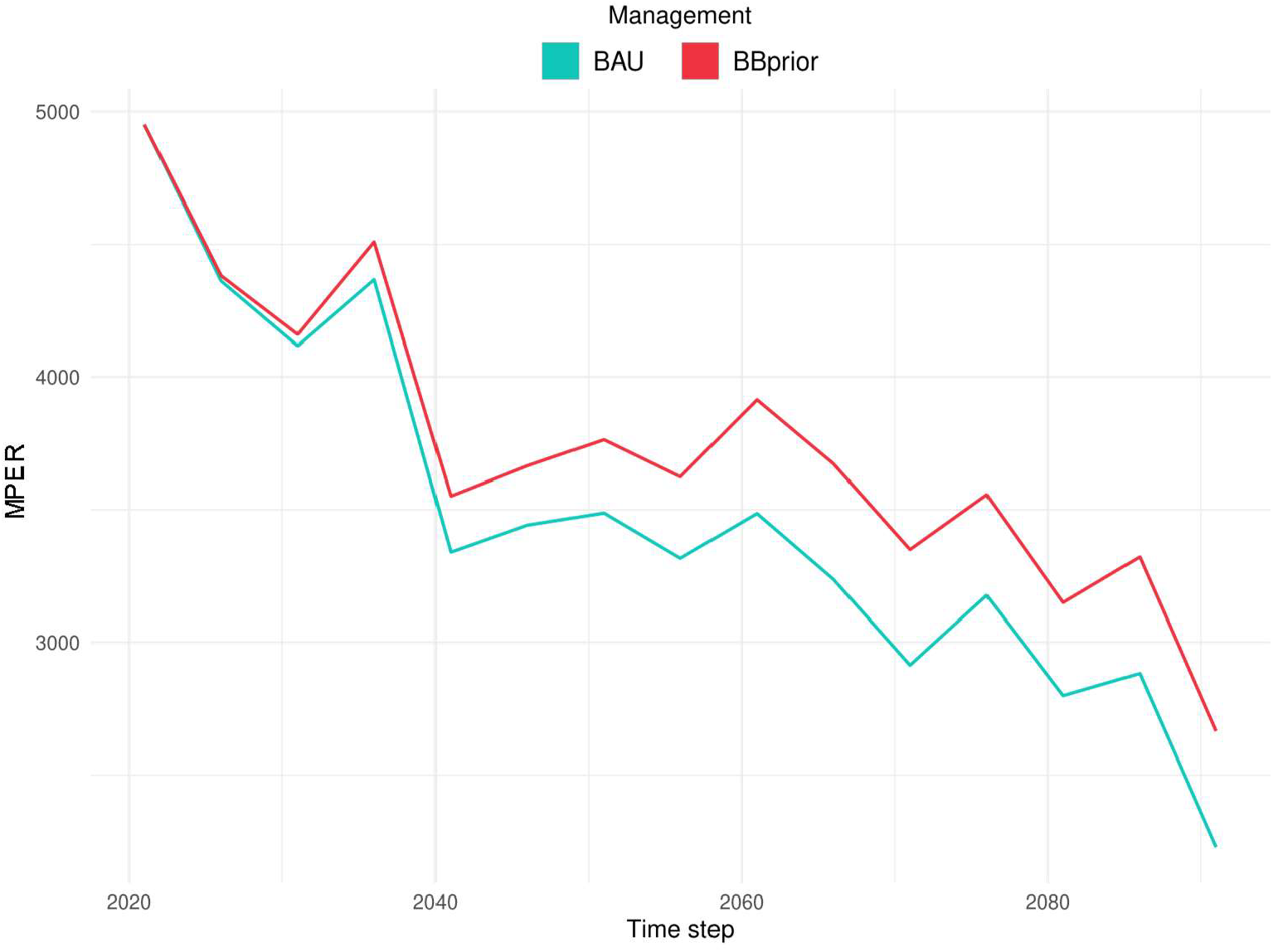
Temporal development of connectivity between forest stands with high susceptibility to spruce bark beetle damage under the Representative Concentration Pathway 4.5 (RCP4.5) climate trajectory. Connectivity was estimated using the mean pairwise effective resistance (MPER), where lower MPER values reflect more potential pathways for movement and higher overall connectivity. MPER was estimated for two forest management regimes: business as usual (BAU) and prioritize harvesting in stands with high susceptibility to damage (BBprior).

**Figure 5.**
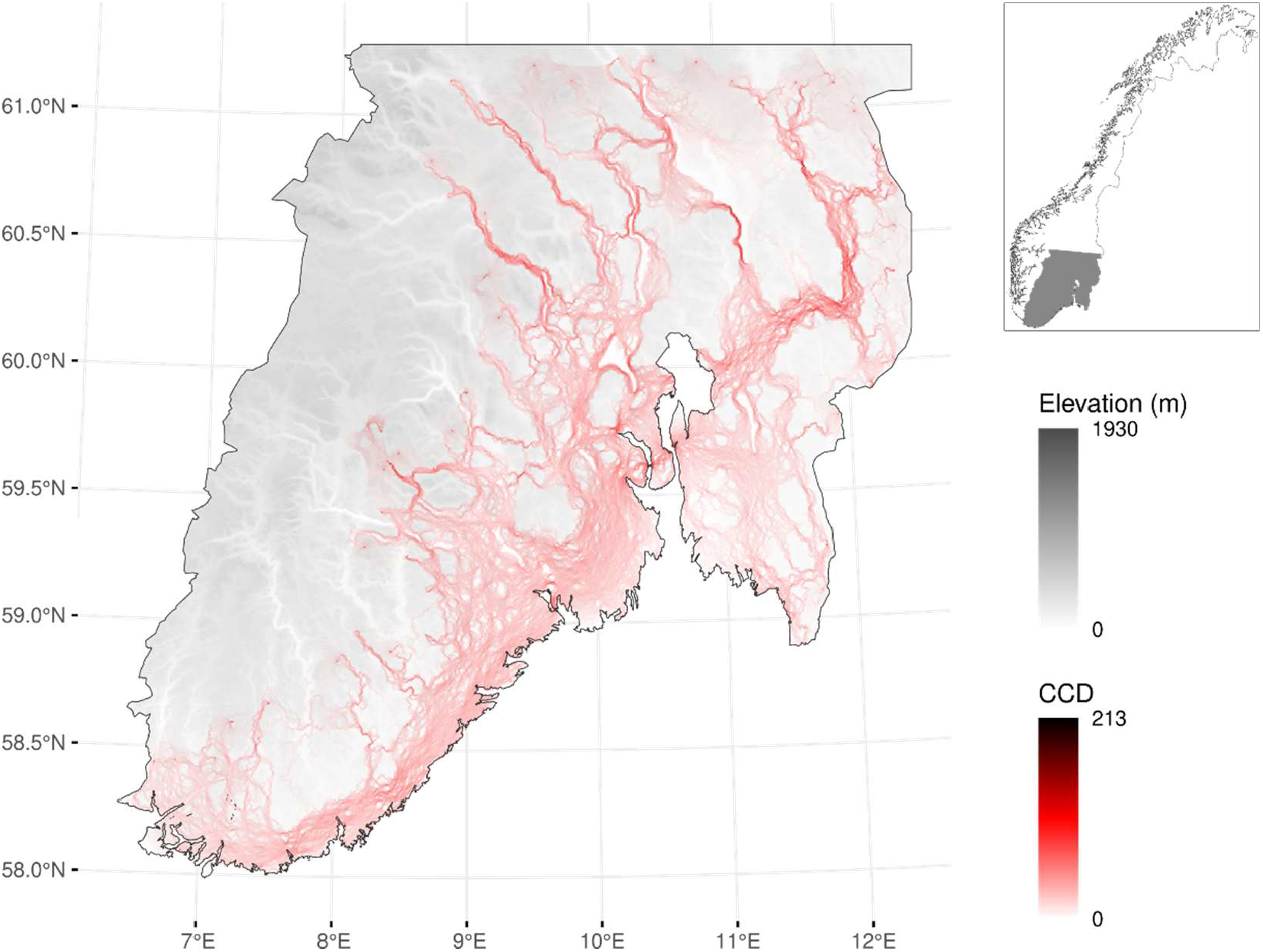
Cumulative current density (CCD) map produced by Circuitscape during the connectivity modeling for the Representative Concentration Pathway 4.5 (RCP4.5) climate trajectory, the prioritize harvesting in stands with high susceptibility to damage (BBprior) management regime, and simulation timestep 2. The map visualizes current flows across the landscape (net movement probabilities), highlighting connectivity patterns that can help identify landscape corridors or pinch points where dispersers are most likely to pass. The inset shows the location of the study area in Norway.

## Discussion

This study examined how forest susceptibility to spruce bark beetle (SBB) damage is shaped by two contrasting management strategies: a conventional business-as-usual (BAU) regime and an approach that prioritizes harvesting in stands with high susceptibility. Through high-resolution simulations covering the full extent of Norway’s forested landscape, we explored forest responses under different climate trajectories over an extended planning horizon (2021– 2100). By integrating a stand-level susceptibility index with scenario analysis and spatial connectivity metrics, our findings highlight how incorporating risk-related information into harvest decisions can modify long-term vulnerability while maintaining national timber supply targets.

While our study focuses on the inherent susceptibility of forests to bark beetle damage, it does not aim to predict actual outbreak dynamics or damage occurrence (Nordkvist et al., 2023). Unfortunately, Norway lacks a consistent national dataset documenting bark beetle infestation rates or associated losses (Machado Nunes Romeiro et al., 2024). Developing robust disturbance models calibrated with local data remains a long-term research objective, likely requiring decades of observations before becoming operationally useful. In the absence of such models, we adopted an alternative approach: assessing how forest management decisions influence the underlying predisposition to damage. This susceptibility-based framework, combined with scenario-based simulations, is already being applied in other countries such as Sweden and Switzerland, where indicator-based assessments are integrated into decision-support systems to support long-term adaptive management under climate change (e.g., López-Andújar Fustel et al. 2024; Mutterer et al. 2025). In our case, it allows for the exploration of the actionable space available to decision-makers when risk-informed harvest strategies are used to reshape future forest vulnerability.

Even if species mixing has been shown to reduce host availability and increase the presence of natural enemies, both of which contribute to lower outbreak potential (Klapwijk et al., 2016), we chose to analyze only stands where Norway spruce comprises more than 50% of the tree cover. It could be reasonably argued that this decision excludes many low-risk forest types, particularly broadleaf-dominated and mixed conifer stands where spruce is only a minor component. However, our decision was intentional, as it allowed us to focus on forest types that are structurally and compositionally most relevant to SBB outbreaks. This focus is further justified by the fact that Norway spruce accounts for approximately 70% of the annual harvest volume in Norway (Breidenbach, Granhus, et al., 2020). While the >50% threshold does not exclude all mixed stands, it provides a sufficiently inclusive yet targeted definition, allowing for the inclusion of structurally mixed forests where spruce is still ecologically significant. Moreover, given the formulation of the SBBSI used here, we a priori expect susceptibility to be generally low in stands where spruce is not the dominant species. Limiting the analysis to spruce-dominated areas therefore provides a clearer signal of how management decisions influence susceptibility. We point out, however, that in all scenarios simulated here, no species change occurs, as stands are replanted with the same species after harvesting. Future research could focus on alternative management strategies where species composition is actively modified, particularly toward increasing broadleaf presence.

The largest differences in susceptibility values were driven by the management regimes, with climate change increasing absolute index values across all regimes but not substantially altering the relative differences established among them. In general, any management strategy that involved forest harvesting was able to reduce national-level susceptibility compared to a scenario where all stands were set-aside from management (NoH). Similar patterns have been reported by Mutterer et al. (2025), who found that increasing management intensity generally leads to lower stand-level predisposition to disturbance. However, while such strategies may enhance disturbance resistance, they may not support other forest values to the same extent. For example, recent evidence indicates that setting aside forest land can deliver greater biodiversity conservation benefits than diversified management alone (Duflot et al., 2022). These contrasting outcomes underscore potential trade-offs: prioritizing disturbance mitigation through active management may come at the cost of other objectives, such as biodiversity conservation or broader ecosystem multifunctionality (Mutterer et al., 2025). Although disturbances are often associated with negative impacts on ecosystem services (Moos et al. 2023), they can also generate positive effects on biodiversity, as evidenced by disturbance-induced shifts in bird community dynamics (Rey et al., 2019; Wermelinger et al., 2025). In this context, management alternatives based on close-to-nature forestry principles (promoting stand structural heterogeneity, site-adapted species, and natural regeneration) have proven particularly effective in enhancing forest resistance to bark beetle disturbances (Mohr et al., 2024) while also supporting the sustained provision of ecosystem services (Blattert et al. 2024). However, as noted by Mutterer et al. (2025), in the context of accelerating climate change impacts, management strategies may need to prioritize reducing disturbance predisposition, even if this entails trade-offs with the provision of specific biodiversity and ecosystem services. Therefore, careful planning and integrated land-use strategies will be essential to minimize these trade-offs and balance multiple forest management objectives. BAU and NoH showed very similar trajectories in terms of both mean SBBSI and the extent of risk-free area, with only marginal differences in favor of BAU. As previously noted, the BAU regime involves relatively low harvesting rates, with only about 0.45% of the forest area in Norway harvested annually. Although the cumulative effect of such interventions may become more noticeable over the full simulation period, the annual impact on forest conditions remains relatively limited. As a result, the capacity of BAU to substantially alter stand susceptibility at the national level is only slightly greater than in a no-harvest scenario. Moreover, commercially mature stands typically exhibit large volumes and tree diameters, which also correspond to higher susceptibility to bark beetle attacks. Therefore, harvesting such stands, as is done under BAU, can contribute to a modest reduction in overall susceptibility. However, since harvest allocation in BAU is primarily driven by economic factors, including stand profitability and accessibility, rather than risk indicators, its impact in reducing bark beetle susceptibility is ultimately limited. In practice, this results in a management outcome that, in terms of susceptibility reduction, is functionally very similar to a no-management scenario. In contrast, the BBprior regime, though not intended as a realistic or prescriptive management plan, serves as a conceptual benchmark. By selecting stands only based on their susceptibility rankings, it isolates the effect of spatially informed harvest allocation. This provides a practical estimate of the maximum reduction in susceptibility that can be achieved when decisions are driven exclusively by risk indicators. Around the year 2036, when the divergence among strategies was greatest, BBprior reduced national-level susceptibility by approximately 5.6% relative to the initial value, while BAU and NoH increased by about 4.1%. By the end of the simulation, BBprior remained 3.7% below baseline, whereas BAU and NoH increased by nearly 10%, resulting in a relative difference of over 13% between approaches (Fig. 1). Nevertheless, mean susceptibility values tend to plateau toward the end of the simulation across all management strategies, suggesting that these represent the upper bounds of what could be achieved through the tested management approaches. This is particularly relevant considering that, by the end of the simulated period, the proportion of spruce-dominated stands classified as risk-free under the BBprior scenario reached approximately 99.1%. Thus, even under an extreme strategy like BBprior, where harvesting is fully guided by susceptibility rankings, it becomes evident that the capacity to further reduce national-level susceptibility diminishes over time. This plateau likely reflects the long-term structural consequences of even-aged management, which leads to increased age-class homogeneity. Such homogenization reduces the flexibility to continue prioritizing the most susceptible stands, while maintaining harvest levels that meet timber demand.

Beyond average susceptibility values, the distribution of standing spruce volume across SBBSI classes reveals important differences in how management strategies influence forest vulnerability over time (Fig. 3). Under BBprior, total spruce volume declines progressively as harvesting consistently removes the most susceptible stands, shifting the remaining biomass toward lower SBBSI classes. This approach not only reduces mean susceptibility but also limits the amount of timber concentrated in highly vulnerable conditions. In contrast, NoH allows susceptible volume to accumulate, resulting in a growing concentration of biomass in the highest SBBSI classes as unmanaged stands mature. BAU displays an intermediate pattern: while it removes some high-susceptibility volume, particularly from the uppermost classes, it does not prevent the overall build-up of vulnerable biomass, especially under warmer climate conditions. These trends are amplified under the RCP4.5 scenario, which accelerates susceptibility transitions and leads to a greater share of standing volume falling into the highest SBBSI categories. By 2091, most spruce volume under RCP4.5 is concentrated in the two most susceptible classes, underscoring the cumulative effect of warming and insufficiently targeted management. In this context, prioritizing the harvest of highly susceptible stands may offer a practical means of redistributing structural vulnerability across the landscape and reducing the likelihood of preventable timber losses.

Spatial maps of SBBSI (Figure 2) provided insights into the geographic distribution of susceptibility under the BBprior regime and the RCP4.5 climate trajectory. Over time, high-susceptibility areas became increasingly fragmented and less prevalent, a direct consequence of harvesting that prioritized the most vulnerable stands. This targeted approach proved particularly effective in southern Norway, where the largest contiguous spruce-dominated forests are located. By 2019, high-SBBSI patches had been substantially reduced, demonstrating the potential of susceptibility-guided harvesting to manage risk at the landscape scale. Importantly, beyond changes in average susceptibility, the spatial configuration of high-risk areas also plays a critical role in outbreak dynamics. Our connectivity analysis revealed that forest management regimes strongly influence the spatial cohesion of susceptible patches. The BBprior strategy consistently produced higher mean pairwise effective resistance (MPER) values than BAU, indicating lower connectivity between highly susceptible stands. This suggests that risk-informed harvesting not only reduces the amount of vulnerable forest but also disrupts the continuity of those areas, potentially constraining the spread of bark beetle infestations. Over time, MPER declined under all regimes, reflecting the landscape-level homogenization associated with even-aged forestry. However, BBprior consistently mitigated this decline, preserving spatial resistance between vulnerable patches to a greater extent. These results highlight the added value of integrating spatial metrics such as MPER into forest planning, as they offer complementary insights into how different management strategies shape not just the magnitude but also the spatial distribution and connectivity of risk.

Although the BBprior strategy demonstrates that spatially targeted harvesting based solely on susceptibility values can significantly reduce average forest risk and fragment high-susceptibility zones, our results also reveal its limitations in the long term. Toward the end of the simulation period, we observed a slight increase in mean susceptibility values (Figure 1) alongside a reduction in landscape resistance (Figure 4), reflecting a broader trend of structural homogenization caused by even-aged management. This suggests that, while spatial prioritization can deliver early gains, it may be insufficient to maintain low susceptibility levels over time unless combined with more fundamental changes to stand structure or regeneration strategies as noted by Muttered et al. (2024) and Seidl et al. (2018). Importantly, while our use of cumulative current density (CCD) maps was illustrative (Fig. 5), these spatial outputs hold valuable potential for future applications. By highlighting landscape-level movement probabilities and identifying key connectivity features, such as corridors and pinch points, CCD data could help guide more nuanced forest management strategies aimed at disrupting the spatial continuity of high-risk zones. Future research could leverage this information to design risk mitigation plans that not only reduce average susceptibility but also strategically reshape landscape connectivity in ways that limit the spread potential of bark beetle outbreaks.

## Notes

### Competing Interest Statement

The authors have declared no competing interest.

## References

Anantharaman, R., Hall, K., Shah, V. B., & Edelman, A. (2020). Circuitscape in Julia: High Performance Connectivity Modelling to Support Conservation Decisions. Proceedings of the JuliaCon Conferences, 1(1), 58. 10.21105/jcon.00058

Anderegg, W. R. L., Hicke, J. A., Fisher, R. A., Allen, C. D., Aukema, J., Bentz, B., Hood, S., Lichstein, J. W., Macalady, A. K., McDowell, N., Pan, Y., Raffa, K., Sala, A., Shaw, J. D., Stephenson, N. L., Tague, C., & Zeppel, M. (2015). Tree mortality from drought, insects, and their interactions in a changing climate. New Phytologist, 208(3), 674–683. 10.1111/nph.13477

Antón-Fernández, C., Mola-Yudego, B., Dalsgaard, L., & Astrup, R. (2016). Climate-sensitive site index models for Norway. Canadian Journal of Forest Research, 46(6), 794–803. 10.1139/cjfr-2015-0155

Astrup, R., Rahlf, J., Bjørkelo, K., Debella-Gilo, M., Gjertsen, A.-K., & Breidenbach, J. (2019). Forest information at multiple scales: development, evaluation and application of the Norwegian forest resources map SR16. Scandinavian Journal of Forest Research, 34(6), 484–496. 10.1080/02827581.2019.1588989

Beudert, B., Bässler, C., Thorn, S., Noss, R., Schröder, B., Dieffenbach-Fries, H., Foullois, N., & Müller, J. (2015). Bark Beetles Increase Biodiversity While Maintaining Drinking Water Quality. Conservation Letters, 8(4), 272–281. 10.1111/conl.12153

Blattert, C., Mönkkönen, M., Burgas, D., Di Fulvio, F., Toraño Caicoya, A., Vergarechea, M., Klein, J., Hartikainen, M., Antón-Fernández, C., Astrup, R., Emmerich, M., Forsell, N., Lukkarinen, J., Lundström, J., Pitzén, S., Poschenrieder, W., Primmer, E., Snäll, T., & Eyvindson, K. (2023). Climate targets in European timber-producing countries conflict with goals on forest ecosystem services and biodiversity. Communications Earth & Environment, 4(1), 119. 10.1038/s43247-023-00771-z

Breidenbach, J., Granhus, A., Hylen, G., Eriksen, R., & Astrup, R. (2020). A century of National Forest Inventory in Norway – informing past, present, and future decisions. Forest Ecosystems, 7(1), 46. 10.1186/s40663-020-00261-0

Breidenbach, J., Waser, L. T., Debella-Gilo, M., Schumacher, J., Rahlf, J., Hauglin, M., Puliti, S., & Astrup, R. (2020). National mapping and estimation of forest area by dominant tree species using Sentinel-2 data. Canadian Journal of Forest Research, 51(3), 365–379. 10.1139/cjfr-2020-0170

Bright, R. M., Cattaneo, N., Antón-Fernández, C., Eisner, S., & Astrup, R. (2024). Relevance of surface albedo to forestry policy in high latitude and altitude regions may be overvalued. Environmental Research Letters, 19(9). 10.1088/1748-9326/ad657e

Cattaneo, N., Astrup, R., & Antón-Fernández, C. (2024). PixSim: Enhancing high-resolution large-scale forest simulations. Software Impacts, 21, 100695. 10.1016/j.simpa.2024.100695

Duflot, R., Eyvindson, K., & Mönkkönen, M. (2022). Management diversification increases habitat availability for multiple biodiversity indicator species in production forests. Landscape Ecology, 37(2), 443–459. 10.1007/s10980-021-01375-8

Hauglin, M., Rahlf, J., Schumacher, J., Astrup, R., & Breidenbach, J. (2021). Large scale mapping of forest attributes using heterogeneous sets of airborne laser scanning and National Forest Inventory data. Forest Ecosystems, 8, 65. 10.1186/s40663-021-00338-4

Hlásny, T., Krokene, P., Liebhold, A., Montagné-Huck, C., Müller, J., Qin, H., Raffa, K., Schelhaas, M.-J., Seidl, R., Svoboda, M., & Viiri, H. (2019). Living with bark beetles: impacts, outlook and management options. From Science to Policy 8. (European Forest Institute, Ed.). European Forest Institute. 10.36333/fs08

Hoogstra-Klein, M. A., Hengeveld, G. M., & de Jong, R. (2017). Analysing scenario approaches for forest management — One decade of experiences in Europe. Forest Policy and Economics, 85, 222–234. 10.1016/j.forpol.2016.10.002

Huang, S., Eisner, S., Wong, W. K., & Cattaneo, N. (2025). The potential impacts of climate and forest changes on streamflow for micro-, meso- and macro-scale catchments in Norway. Journal of Hydrology: Regional Studies, 57, 102147. 10.1016/j.ejrh.2024.102147

IPCC. (2021). Summary for Policymakers. In V. Masson-Delmotte, P. Zhai, A. Pirani, S.L. Connors, C. Péan, S. Berger, N. Caud, Y. Chen, L. Goldfarb, M.I. Gomis, M. Huang, K. Leitzell, E. Lonnoy, J.B.R. Matthews, T.K. Maycock, T. Waterfield, O. Yelekçi, R. Yu, & B. Zhou (Eds.), Climate Change 2021: The Physical Science Basis. Contribution of Working Group I to the Sixth Assessment Report of the Intergovernmental Panel on Climate Change (pp. 3–32). Cambridge University Press, Cambridge, United Kingdom and New York, NY, USA. https://doi.org/DOI: 10.1017/9781009157896.001

Kautz, M., Dworschak, K., Gruppe, A., & Schopf, R. (2011). Quantifying spatio-temporal dispersion of bark beetle infestations in epidemic and non-epidemic conditions. Forest Ecology and Management, 262(4), 598–608. 10.1016/j.foreco.2011.04.023

Klapwijk, M. J., Bylund, H., Schroeder, M., & Björkman, C. (2016). Forest management and natural biocontrol of insect pests. Forestry: An International Journal of Forest Research, 89(3), 253–262. 10.1093/forestry/cpw019

Landbruksdirektoratet. (2023). Kartlegging av foryngelse og miljøhensyn ved hogst 2022. https://www.landbruksdirektoratet.no/nb/nyhetsrom/rapporter/kartlegging-av-foryngelse-og-miljohensyn-ved-hogst-2022

Lauri, P., Forsell, N., Di Fulvio, F., Snäll, T., & Havlik, P. (2021). Material substitution between coniferous, non-coniferous and recycled biomass – Impacts on forest industry raw material use and regional competitiveness. Forest Policy and Economics, 132, 102588. 10.1016/j.forpol.2021.102588

López-Andújar Fustel, T., Öhman, K., Klapwijk, M., Nordkvist, M., Sängstuvall, L., Lämås, T., & Eggers, J. (2024). Impact of management strategies on forest susceptibility to spruce bark beetle damage and potential trade-offs with timber production and biodiversity. Forest Ecology and Management, 563, 121964. 10.1016/j.foreco.2024.121964

Machado Nunes Romeiro, J., Jostein, G., Paal, K., Tron, E., & and Antón Fernández, C. (2024). Bark beetle damage in Norwegian forests: a study of model suitability and projected impact under climate change. Scandinavian Journal of Forest Research, 39(1), 30–43. 10.1080/02827581.2023.2289648

Maleki, K., Astrup, R., Kuehne, C., McLean, J. P., & Antón-Fernández, C. (2022). Stand-level growth models for long-term projections of the main species groups in Norway. Scandinavian Journal of Forest Research, 37(2), 130–143. 10.1080/02827581.2022.2056632

McRae, B. H., Dickson, B. G., Keitt, T. H., & Shah, V. B. (2008). Using circuit theory to model connectivity in ecology, evolution, and conservation. Ecology, 89, 2712–2724.

Mikkelson, K. M., Dickenson, E. R. V, Maxwell, R. M., McCray, J. E., & Sharp, J. O. (2013). Water-quality impacts from climate-induced forest die-of. Nature Climate Change, 3(3), 218–222. 10.1038/nclimate1724

Müller, M., Olsson, P.-O., Eklundh, L., Jamali, S., & Ardö, J. (2022). Features predisposing forest to bark beetle outbreaks and their dynamics during drought. Forest Ecology and Management, 523, 120480. 10.1016/j.foreco.2022.120480

Mutterer, S., Blattert, C., Bont, L. G., Griess, V. C., & Schweier, J. (2025). Beetles, wind, and fire: Effects of climate change and close-to-nature forestry on disturbance predisposition and ecosystem service trade-offs. Forest Ecology and Management, 586, 122690. 10.1016/j.foreco.2025.122690

Nordkvist, M., Eggers, J., Fustel, T. L.-A., & Klapwijk, M. J. (2023). Development and implementation of a spruce bark beetle susceptibility index: A framework to compare bark beetle susceptibility on stand level. Trees, Forests and People, 11, 100364. 10.1016/j.tfp.2022.100364

Norwegian Ministry of Climate and Environment. (2020). National Forestry Accounting Plan for Norway, including forest reference level for the first commitment period 2021-2025. (pp. 0– 49).

O’Brien, P., Carr, N., & Bowman, J. (2023). Using sentinel nodes to evaluate changing connectivity in a protected area network. PeerJ, 11:e16333. 10.7717/peerj.16333

Seidl, R., Müller, J., Hothorn, T., Bässler, C., Heurich, M., & Kautz, M. (2016). Small beetle, large-scale drivers: how regional and landscape factors affect outbreaks of the European spruce bark beetle. Journal of Applied Ecology, 53(2), 530–540. 10.1111/1365-2664.12540

Sevillano, I., Antón-Fernández, C., Søgaard, G., & Astrup, R. (2025). Improved forest management for increased carbon sequestration: An assessment of the most prominent approaches in Norway. Journal of Environmental Management, 375, 124333. 10.1016/j.jenvman.2025.124333

Tveite, B., & Braastad, H. (1981). Bonitering av gran, furu og bjork. Technical Report 4.

Vergarechea, M., Astrup, R., Fischer, C., Øistad, K., Blattert, C., Hartikainen, M., Eyvindson, K., Di Fulvio, F., Forsell, N., Burgas, D., Toraño-Caicoya, A., Mönkkönen, M., & Antón-Fernández, C. (2023). Future wood demands and ecosystem services trade-offs: A policy analysis in Norway. Forest Policy and Economics, 147, 102899. 10.1016/j.forpol.2022.102899

Wong, W. K., Haddeland, I., Lawrence, D., & Beldring, S. (2016). Gridded 1 x 1 km climate and hydrological projections for Norway. http://hdl.handle.net/11250/2500569

Wulff, S., & Roberge, C. (2022). Nationell riktad Skogsskadeinventering (NRS) -Inventering av granbarkborreangrepp i götaland och Svealand 2022.

Zimová, S., Dobor, L., Hlásny, T., Rammer, W., & Seidl, R. (2020). Reducing rotation age to address increasing disturbances in Central Europe: Potential and limitations. Forest Ecology and Management, 475, 118408. 10.1016/j.foreco.2020.118408

