## Appendix for "A high-resolution, country-wide scenario analysis of forest susceptibility to spruce bark beetle damage in Norway"

**Spruce Bark Beetle Susceptibility Index and Variable calculations**

Volume of Spruce at the stand level

The volume of Spruce at the stand level was calculated by summing the volume (V, $m^3$/ha) of Spruce-dominated pixels within each stand. This sum was then divided by the total number of pixels in the stand, including non-Spruce pixels, to obtain the average Spruce volume at the stand level. Finally, these volumes were converted into relative scores as defined by (Nordkvist et al., 2023).

Mean diameter

This variable is not included in SR16. To address this, we developed an empirical model to predict the basal area-weighted mean diameter (Dgv, cm) using dominant height (H, m) and site index (SI, m) as predictors. The model was fitted using data from 2045 Spruce-dominated plots in the Norwegian National Forest Inventory (NFI). A plot was classified as Spruce-dominated if at least 75% of its timber volume over bark was attributed to Norway Spruce. The Dgv values for each NFI plot were calculated based on the description provided in the Heureka Dictionary (Heureka SLU, n.d.). Dominant height and site index are two variables consistently measured in the NFI. Using the fitted model, we then predicted Dgv for each pixel in SR16 by using the SR16 estimates for H and SI.

The final model, along with its parameters, is as follows:

$$
\begin{aligned}
Dgv_{cm} = \exp\big(&0.93734002 + 0.88401721 \cdot \ln(H_m) + (SI_m = 8) \cdot (-0.05699199) \\
&+ (SI_m = 11) \cdot (-0.19121368) + (SI_m = 14) \cdot (-0.27457949) \\
&+ (SI_m = 17) \cdot (-0.31593645) + (SI_m = 20) \cdot (-0.34632881) \\
&+ (SI_m = 23) \cdot (-0.35552196) + (SI_m = 26) \cdot (-0.32402875)\big) \\
&\cdot 1.023775
\end{aligned}
$$

Temperature sums

To calculate this variable, we used daily mean temperature data from 2016 to 2100. The steps to calculate the temperature sum at the start of each 5-year time step are as follows:

Using the example of 2021 (time step 2021–2026), we subtracted 5°C from the daily mean temperature values between April 1 and August 31 for each year from 2016 to 2020. The threshold of 5°C represents the minimum temperature for degree day accumulation. We then summed these adjusted daily values to calculate the degree days for each year, resulting in five values (one for each year from 2016 to 2020). These were averaged to produce a 5-year mean temperature sum (in degree days) for April 1 to August 31, which was used to compute the SBBSI index for 2021.

In the calculation of the SBBSI for the 2026–2030 time-step, the same process was applied using temperature data from 2021 to 2025. A similar process was repeated for all other time steps in the simulations.

Volume of birch at the stand level

The volume of Birch at the stand level was calculated by summing the volume (V, $m^3$/ha) of Birch-dominated pixels within each stand. This sum was then divided by the total number of pixels in the stand, including non-Birch pixels, to obtain the average Birch volume at the stand level. Finally, these volumes were converted into relative scores as defined by (Nordkvist et al., 2023).

Soil moisture

Soil moisture is included in the SBBSI as a categorical variable with five classes: Dry, Mesic, Mesic-Moist, Moist, and Wet. Since this variable is not available in SR16, we derived its values using the Depth to Water (DTW) map available at NIBIO. DTW is a raster map with a 1-meter resolution that covers the entire surface of Norway. It indicates the vertical distance to the groundwater table. The DTW map is classified as follows:

Class 1: Water

Class 2: DTW from 0 to 25 cm

Class 3: DTW from 25 to 50 cm

Class 4: DTW from 50 to 75 cm

Class 5: DTW from 75 to 100 cm

Class 6: DTW greater than 100 cm

To use this data, we first performed an upscaling of the DTW map to a resolution of 16 meters, assigning the most frequent value within each 16-meter pixel. We then selected the pixels overlapping with the forested areas represented in the SR16 map. Finally, we reclassified the DTW values into soil moisture classes using the following rules:

DTW Class 1 → Soil Moisture Class: Wet

DTW Class 2 → Soil Moisture Class: Wet

DTW Class 3 → Soil Moisture Class: Moist

DTW Class 4 → Soil Moisture Class: Mesic-Moist

DTW Class 5 → Soil Moisture Class: Mesic

DTW Class 6 → Soil Moisture Class: Dry

Soil moisture classes were converted into relative scores as defined by (Nordkvist et al., 2023).

Stand density

To calculate stand density on each pixel, tree heights are classified into intervals (e.g., 0–2 m, 2–4 m). For each height class and site index combination, the 95th and 5th percentiles of stand volume (V, $m^3$/ha) are calculated as maximum ($Q_{95}$) and minimum ($Q_{05}$) reference values. These values define the potential range of stand volume per site quality and height class. Subsequently, stand density is normalized for each pixel using the formula: Density = ($V_{pixel}$ - $Q_{05}$) / ($Q_{95}$ - $Q_{05}$). ). Finally, density values were converted into relative scores as defined by (Nordkvist et al., 2023).

Age structure

Pixels within each forest stand are grouped into age intervals. For each stand, the total stand volume (V, $m^3$/ha) is calculated, and the proportion of this volume contributed by each age class is determined. A stand is classified as “even-aged” if a single 5-year age class contributes at least 95% of the total stand volume. A stand is considered “mostly even-aged” if a single 20-year age class accounts for at least 80% of the total volume. Finally, stands not meeting these criteria are classified as uneven-aged. Age structure classes were converted into relative scores as defined by (Nordkvist et al., 2023).

Index calculation

The index calculation followed the methodology of Nordkvist et al. (2023). For each pixel, if the volume of spruce was > 0, the Dgv of spruce > 20 cm, and the temperature sum ≥ 745, the relative scores of the indicator variables were multiplied by their corresponding weightings to obtain the final values. These values were then summed to compute the SBBSI index for each pixel.

**References**


**Heureka SLU**. (n.d.). *Dictionary*. Retrieved December 27, 2024, from https://www.heurekaslu.se/wiki/Dictionary#Dgv_-_Basal_area_weighted_mean_diameter

**Nordkvist, M.**, Eggers, J., Fustel, T. L.-A., & Klapwijk, M. J. (2023). Development and implementation of a spruce bark beetle susceptibility index: A framework to compare bark beetle susceptibility on stand level. *Trees, Forests and People*, *11*, 100364. https://doi.org/https://doi.org/10.1016/j.tfp.2022.100364
